# Neural systems underlying the learning of cognitive effort costs

**DOI:** 10.1101/2020.06.08.139618

**Authors:** Ceyda Sayali, David Badre

## Abstract

People balance the benefits of cognitive work against the costs of cognitive effort. Models that incorporate prospective estimates of the costs of cognitive effort into decision making require a mechanism by which these costs are learned. However, it remains open what brain systems are important for this learning, particularly when learning is not tied explicitly to a decision about what task to perform. In this fMRI experiment, we parametrically manipulated the level of effort a task requires by increasing task switching frequency across six task contexts. In a scanned learning phase, participants implicitly learned about the task switching frequency in each context. In a subsequent test phase, participants made selections between pairs of these task contexts. We modeled learning within a reinforcement learning framework, and found that effort expectations that derived from task-switching probability and response time (RT) during learning were the best predictors of later choice behavior. Prediction errors (PE) from these two models were associated with FPN during distinct learning epochs. Specifically, PE derived from expected RT was most correlated with the fronto-parietal network early in learning, whereas PE derived from expected task switching frequency was correlated with the fronto-parietal network late in learning. These results suggest that multiple task-related factors are tracked by the brain while performing a task that can drive subsequent estimates of effort costs.

## Introduction

People avoid cognitively effortful tasks when given the option (Kool et al., 2010; Westbrook et al., 2013). Based on this observation, it has been proposed that effortful tasks incur a subjective cost that drives demand avoidance. Further, in order to weigh the expected effort cost of a task against its expected rewards, people must learn to predict the level of cognitive effort required to successfully perform a task (Shenhav et al., 2013; Botvinick, 2007).

In the laboratory, demand avoidance has been tested using effort discounting paradigms. In these tasks, participants learn the association between a unique task identifier and a level of subjective task difficulty during a learning phase. In the following decision phase, participants engage in repeated decision-making between an easy task paired with smaller reward or a difficult task paired with higher reward. Several fMRI studies showed that people discount the value of an offer as a function of their subjective cost during the decision phase (Massar et al., 2015; Chong et al., 2017; Westbrook et al., 2019; 2020).

As decision-making involves comparing the costs and benefits of a given action, it has been proposed that diverse factors related to value are integrated into a unitary or common currency subjective value representation (Padoa-Schioppa, 2011). The factors related to the cost side of the equation might include physical costs, time delays, riskiness, as well as the motivational state of the agent, their representation of themselves, and the opportunity costs they perceive. Across economic decision making tasks that have tested a range of these factors, such as delay discounting (Massar et al, 2015), physical effort avoidance (Chong et al., 2017; Croxson et al., 2009) or reinforcement learning (Jocham et al., 2011), a network of brain areas has been consistently observed to track the value of the chosen option (Levy & Glimcher, 2012) with distinct clusters in VMPFC, VS and PCC (Clithero & Rangel, 2014). This domain-general network is called the ‘Subjective Value (SV) Network and has been shown to predict choice behavior as a function of both the costs and benefits of an action. With particular relevance to the present study, a recent fMRI study showed that the same SV network also tracked the subjective value of cognitive effort (Westbrook et al., 2019), while control-related brain regions of the fronto-parietal network (FPN) tracked the decision difficulty between offer options, and not the SV of the task.

These observations were partially consistent with those from a previous fMRI study of effort-based decision making (Sayali & Badre, 2019). In that study, we separated learning and decision-making phases of an effort selection task. Participants first implicitly learned about the cost associated with each of six parametrically increasing effort levels, without being informed about the subsequent decision phase. During the subsequent decision phase, they selected which task to perform based on this prior experience. As with Westbrook et al. (2019), we found that during the decision phase, though FPN activity tracked task difficulty, it was not related to effort-based predictions. In our study, the only predictor of effort avoidance during task performance was default mode network activity (DMN). Notably, this network includes the vmPFC which overlaps the SV network highlighted by Westbrook et al. (2019). Though, we did not find evidence of a relationship between other nodes of the SV network and effort avoidance, including the ventral striatum and dACC. Overall, however, studies of effort-based decisions appear to agree that SV-related regions, like vmPFC, encode effort costs, while other networks involved in cognitive control, like the FPN, track difficulty but not the subjective cost driving avoidance decisions.

Importantly, however, these prior studies have focused on the decision stage, when people are familiar with the tasks involved and are weighing costs and benefits of cognitive effort. Few experiments have tested how the effort costs associated with tasks themselves are acquired. The one fMRI experiment to date that focused on effort learning observed that task difficulty (e.g. task switching) and performance-based prediction errors (e.g. response time) drove the acquisition of effort costs (Nagase et al., 2018). Further, expected costs positively activated dACC, as well as vmPFC, a region in the SV network. However, in this task, effort decisions were made while effort costs were also being learned. This still leaves open the possibility that these regions might be tracking the effort costs only because an explicit decision about task value is due (i.e., learning about the decision), rather than because an implicit cost was being learned. As such, it is not presently known if the SV network is involved when learning effort costs or tracking task value in the absence of a decision about what task to perform.

In order to fill this gap in the literature, we ask what neural mechanisms underlie implicit effort learning in the absence of decision making. We use the parametric effort selection task from Sayali & Badre (2019). We test the involvement of the brain networks observed during effort-based decision-making (FPN, DMN in Sayali & Badre, 2019; SV regions in Westbrook et al., 2019) using established learning models of cognitive effort costs, as defined by task-switching and response time (Nagase et al., 2018). Our overall hypothesis is that SV regions will negatively scale with the expected effort costs, and the magnitude of this effect will become stronger in later phases of learning with greater task experience.

## 2. Methods

The current study asks what neural mechanisms underlie the implicit learning of effort costs from task experience, while in the absence of effort-based decisions. We hypothesize that 1) effort costs are learned through experience and predict later effort selections, 2) task difficulty and performance-based prediction errors drive the learning of effort costs, 3) SV brain regions track expected effort costs. In order to test these hypotheses, we estimated expected costs and prediction errors during an implicit learning phase based on variants of the cost prediction error model of effort. We then tested correlates of these parameters using model-based fMRI analysis of the brain during learning.

### 2.1. Participants

Three-hundred and seventeen adults were pre-screened in the lab using the Need for Cognition (NfC) Inventory. The Need for Cognition (NfC) inventory was developed to assess the participant’s tendency to engage in cognitively challenging tasks in their daily lives (Cacioppo & Petty, 1982). In Sayali and Badre (2019), we found considerable individual variability in effort selection behavior when using the Demand Selection Task (DST) in fMRI. Specifically, participants were evenly split between effort avoiders who tended to choose the easier task more frequently and effort seekers who tended to choose the harder task more frequently. This split was notably different from both prior studies using the DST and our own behavioral work with this task.

In order to understand this difference, we conducted a separate study and found that individuals who volunteer for fMRI, in general, tended to score higher on the NfC compared to participants who only volunteered for behavioral studies. This can lead to a self-selection bias in fMRI versions of DST in which participants tend to be more effort seeking than would be encountered in the general population, and this bias threatens representativeness and so generalizability when left uncontrolled. This is important because Sayali and Badre (2019) found that effort seekers differed from effort avoiders in their behavioral and some neural responses. Thus, our screening procedures were designed to control for self-selection by recruiting demand avoiding individuals for this fMRI study. Though, this does mean that our conclusions are intended to generalize to demand avoidant individuals generally.

Given these considerations, we only included participants scoring 16/72 or lower on the NfC inventory (see Supplementary fig. 1 for the distribution of participants). We further screened the remaining 119 participants based on neurological or psychiatric diagnosis, drug use, or contraindication for MRI. 78 participants met the fMRI eligibility criteria, agreed to be re-contacted, and had not participated in a previous study that tested effort-based decision-making. From these, 29 agreed to participate in our fMRI study. However, 7 of these cancelled their appointment on the day of their scan or withdrew. Given that our participants were prescreened as highly demand avoiding, this rate of withdrawal was not unexpected. However, we consider the implications of this sampling procedure for generalizability of this study in the discussion.

**Figure 1.**
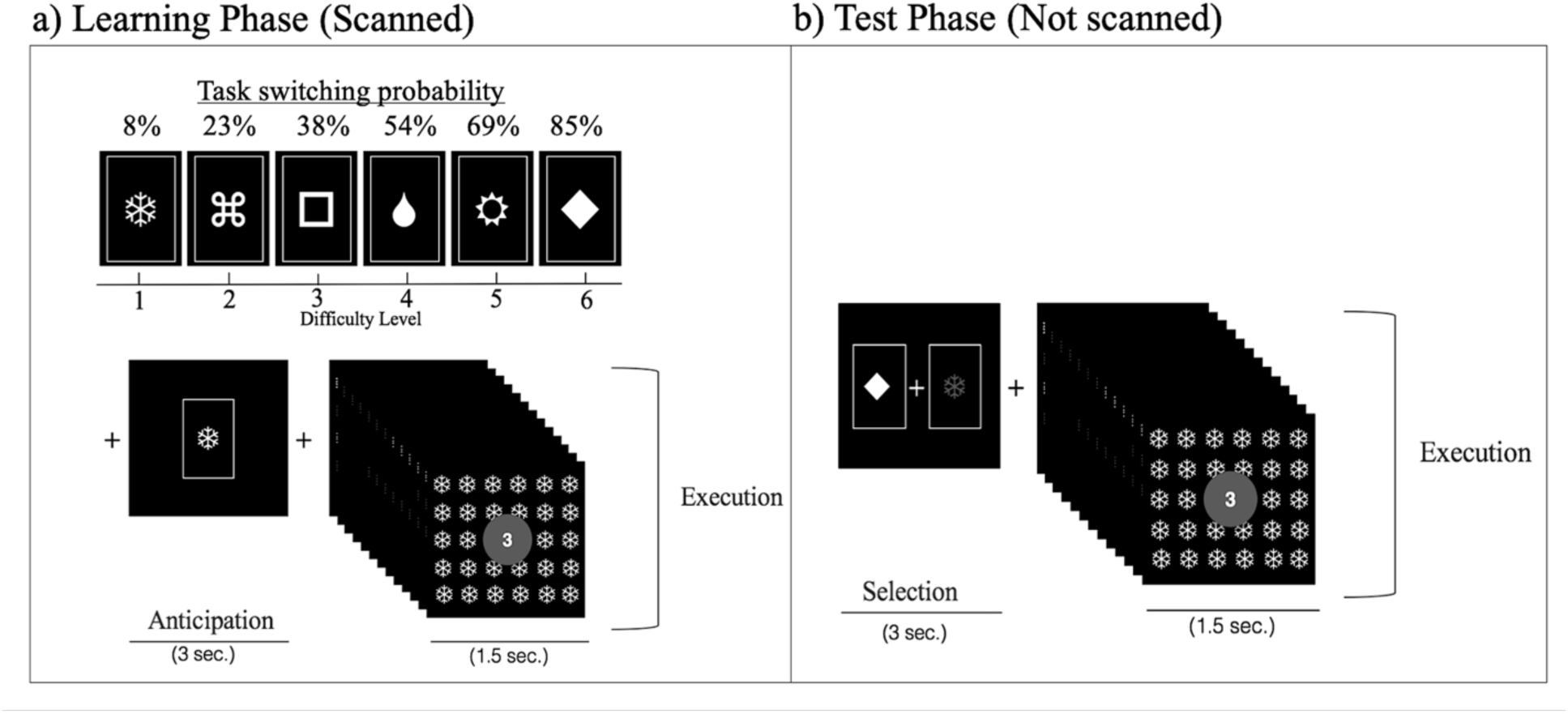
Illustration of the task design. (a) In the Learning Phase, participants learned the association between difficulty levels and corresponding unique task identifiers (top) while being scanned. Anticipation and Execution epochs (bottom) were separately optimized in a hybrid fMRI design. During the Anticipation epoch, participants viewed a symbol icon (a virtual card deck) for 3 seconds. Then, they performed the difficulty level associated with that symbol, while the symbol was tiled on the background. (b) Following the Learning Phase, participants completed the Selection Phase outside the scanner. During a Selection epoch, participants chose between two of the symbols they learned in the Learning Phase. During the Execution epoch, they performed the difficulty level that was associated with their choice.

Therefore, a total of 22 right-handed adults (median age = 20 [18-34]; 12 female) who scored lower than 16/72 on NfC, with normal or corrected to-normal vision were scanned in the fMRI experiment. 3 additional participants were excluded prior to analysis because they either failed to follow instructions (2 participants) or moved more than a voxel in all sessions (1 participant). Thus, a total of 19 participants were included in the behavioral and fMRI analyses. Participants provided informed consent in accordance with the Research Protections Office at Brown University. All participants were compensated for their time either monetarily or with course credit.

### 2.2. Overview of the Behavioral Task

Participants performed a parametric variant of the DST, closely following the procedures from Sayali and Badre (2019; Fig 1). As an overview, the task was performed in two phases. In an initial, scanned Learning phase, the participants performed six tasks and associated each with a virtual “card deck”, denoted by a particular symbol that tiled the screen when that task was performed. The six tasks differed from each other in the proportion of task switching required which manipulated the subject effort associated with each, as described below.

In a second Test phase, participants chose which deck they wished to perform (Selection epoch) and then performed a series of trials drawn from that deck (Execution epoch). We now provide detail on each of these phases of the behavioral task.

#### 2.2.1. The Task “Decks” and Basic Task Run Structure

Throughout all phases of the experiment, participants performed blocks consisting of 13 trials “drawn” from a virtual deck (Fig 1a). On each trial, the participant categorized a centrally presented digit as either odd/even (parity judgment) or greater/less than 5 (magnitude judgment). The color of a circle (green or blue) surrounding the digit cued whether to perform the parity or magnitude judgment on each trial. The participant indicated their categorization by pressing the left or the right arrow key on the keyboard. The response deadline was 1.5 sec. Trials were separated by .2 sec. The mappings of color to task and from each categorization to a left or right arrow press were provided in an instruction before the experiment and were consistent throughout both phases of the experiment. Response mappings and color-to-task mappings were counterbalanced across participants. Digits 1-4 and 6-9 were used with equal frequency across both tasks and were randomized for order of presentation.

In order to manipulate cognitive effort between the decks, we varied the frequency of task switching required by a particular deck (Fig 1a), as in how often a trial-to-trial transition required going from a parity to magnitude judgment or vice versa. We assumed that more task switches in a deck would also require more cognitive control for that deck and so more cognitive effort. Cognitive control selects and guides our actions in accordance with our current goals and the current context (Norman & Shallice, 1986; Miller and Cohen, 2001; Badre, 2020). This process is important when one needs to use the context to enact shifts between multiple tasks that involve different sets of rules that demand distinct behavioral outputs based on the same inputs. Thus, more frequent task switching requires greater cognitive control and has been shown to lead to more effort avoidance (Kool et al., 2010).

As a short hand for this logic, we will refer to the proportion of task switching in a deck as its *effort level*. Importantly, participants were not informed about the effort difference between decks; they had to learn about these differences through experience.

The probability of switching tasks increased across decks over six effort levels: 8%, 23%, 38%, 54%, 69%, 85% (Fig 1a, top). Importantly, in the Learning Phase, we kept the probability of encountering a magnitude and a parity trial constant across execution blocks. In other words, no effort block was associated with a greater likelihood of performing either task. What differed was the transition probability between the two tasks (i.e., the likelihood of switching). Thus, any differences in effort preference among the tasks could not be attributed to differences in the difficulty or effort of the categorizations themselves, but rather were attributable to the task switching manipulation. Further, the lowest effort level still included one switch. We did not include “pure blocks” of only parity judgments or only magnitude judgments with zero switches, as doing so would have made these decks qualitatively different from all other decks, in terms of the amount of each task being performed.

At the beginning of each run, a shape was presented for 3 sec to indicate which deck was about to be performed, and this shape was also tiled in the background throughout the performance of the run. Participants were told that this shape represented the symbol on the back of the virtual card deck from which trials for that sequence were drawn. Thus, each effort level could be associated with a particular deck symbol. Participants could learn this relationship through experience with the tasks. The mappings between deck symbols and effort levels were randomized across participants.

#### 2.2.2. Practice Phase

During an initial “Practice Phase” participants gained familiarity with the trial structure, the categorization decisions, and the color and response mappings. After being given instructions regarding the categorization and response rules, they practiced two 13-trial blocks. During the presentation of these blocks, the researcher closely monitored the performance of the participant and assisted the participant if needed. This phase was repeated if the participant asked for a repeat of the task rules, and thus the data from this phase was not recorded.

Each task sequence included the presentation of a deck symbol (3 s), and the subsequent performance of the categorization trials. In the first run of the Practice Phase, feedback was presented after the button press as either ‘Correct’ in pink, ‘Incorrect’ in yellow, or ‘Please respond faster’ if the participant failed to answer before the deadline (1.5s). In the second run of the Practice phase, feedback was omitted as would be the case in the actual experiment. The deck symbols that participants encountered in the Practice Phase were for practice only and were not presented during the Learning or Test phases to avoid any association of difficulty or error feedback with a particular deck symbol during this phase.

#### 2.2.3. Learning Phase

In the Learning Phase (Fig. 1a), participants learned the association between the six deck symbols and each effort level. Each deck was performed 15 times. The Learning Phase blocks were randomized by using an event-related fMRI optimization tool (optseq2). This method automatically schedules fMRI events by randomizing the task order and jittering the time interval between events. It generates thousands of these orders from which it picks those orderings with the best efficiency. This phase was performed during fMRI scanning, and participants used an MRI-compatible button box to indicate their decisions The Execution trials were optimized as blocks, and not as single-trial events in a hybrid design where anticipation epochs were optimized as single events.

The Anticipation and Execution events were separated in time by a jittered time interval (mean 4 secs) so that signal related to each could be analyzed independently. The Learning phase was separated into two, approximately 20 minute-long scanning runs in order to estimate the effects of early vs. later stage effort learning on brain activity.

#### 2.2.4. Test Phase

In the Test Phase (Fig 1b), two decks were presented and the participant chose which to perform (Selection epoch). The Test Phase took place outside the magnet, in a behavioral testing room immediately following the MRI session. The participants were told to freely choose between decks prior to a 3 sec deadline. We note that in contrast to other DST variants (Gold et al., 2014), participants in this task were not told about the difficulty manipulation to avoid biasing choices based on participants’ explicit beliefs about effort. Once the participant made their selection, the selected deck turned blue and both options remained on the screen until the end of the 3 sec deadline. If the participant missed the deadline, the same choice pair was repeated at the end of the experiment until all selections had been made and executed.

Each choice pair was selected from the set of fifteen unique (un-ordered) pair combinations of all six decks, excluding self-pairing (e.g., deck #1 paired with deck #1). Each deck was presented either on the left or the right side of the screen, counterbalanced for location across trials. The Selection epoch was followed by performance of the selected effort level task deck (Execution epoch). The sequence of events, timing, and response mappings during Execution were the same as during the Learning phase. The Test phase was separated into four, approximately 15 minute-long blocks. The Test Phase trials were pseudo-randomized so that each run contained an equal number of each selection pair. In each run, each pair was presented 3 times, making a total of 180 decision trials across 4 blocks.

#### 2.2.4. Post-experimental questionnaire

Upon the completion of the experiment, participants were asked to answer by free response text entry to 6 questions regarding their understanding of the task. The questions were as follows: 1) What do you think the deck symbols stood for?, 2) Was there any reason why you selected certain decks in the last phase?, 3) Did you have a preference for any deck?, 4) What do you think the experimental manipulation was?, 5) Imagine somebody else is about to perform the same experiment. What advice would you give them? And, did you follow this advice?, 6) Please rate each deck symbol in terms of its difficulty (out of 6). Free responses that included terms such as ‘difficulty’, ‘easy’, ‘hard’ and ‘effort’ were scored as ‘awareness of the effort manipulation’. Accuracy of the last inventory question was determined by calculating the match and mismatch between participants’ ranking of a given symbol with the difficulty it was assigned.

### 2.3. Behavioral Data Analysis

Trials with response times (RT) below 200 ms were excluded from further analysis. Execution trials on which participants missed the response deadline were also excluded (approximately 1% of trials in both phases). Task performance was analyzed with effort level and experimental phases (Learning or Test phase) as within-subject variables. One participant never chose the 5^th^ effort level during the Test Phase, and thus was necessarily excluded from the effort*phase performance analysis (see below).

Choice behavior was assessed by calculating the probability of selecting each effort level across all selection trials on which that effort level was presented as an option. The decision difference analyses calculated the choice probability and the decision time to select the easier task across all decisions with the same difference in difficulty levels between effort options. For example, choice probability associated with a difficulty difference of 1 would be computed by averaging the probability of choosing the easier task across all choice pairs that differed by 1 effort level (i.e., 1 vs 2, 2 vs 3, 3 vs 4, 4 vs 5 and 5 vs 6).

Data were analyzed using a mixed-design analysis of variance (ANOVA) (within subject factor: Effort). If the sphericity assumption was violated, Greenhouse-Geisser correction was used (Greenhouse and Geisser, 1959). Significant interactions were followed by simple effects analysis, the results of which are presented with False Detection Rate (FDR) correction. Alpha level of .05 was used for all analyses. Error bars in all figures stand for within-subject error.

#### 2.3.1. Computational model fitting

To test our hypothesis regarding effort cost learning, we modeled the acquisition of effort associations within a reinforcement learning framework (Sutton and Barto, 2018; Daw, 2011). We specifically used a prediction error (PE)-based model that has been applied during an effort selection task (Nagase et al., 2018). Following this prior work, we tested separate models that learned effort costs from prediction errors in 1) task-switching probability, 2) response time during effort execution, 3) error rates during effort execution. We tested two models that relied on prediction error in response time, as described below. Thus, overall, we tested four models of effort cost learning.

Note that our models do assume that participants did not learn the cost of the effort tasks during the Selection Phase. We make this simplifying assumption because the selection rates across the four blocks of the Selection Phase are stable, suggesting that learning of the effort costs have been mostly finalized in the Learning Phase, or at the very least, change very little. These findings are also consistent with our earlier study (Sayali & Badre, 2019) which showed that effort selection rates were stable across trials in the Selection Phase.

##### 2.3.1.1. Task Switch Cost Model

The Task Switch Cost Model attempts to learn the likelihood of a switch based on the context.

To begin, the model assumes an even likelihood of a switch or repeat and so the expected cost on the first trial was initialized to 0.5. Then, on each effort execution block of the Learning Phase, expected cost of the effort task is updated by a PE of that effort level:

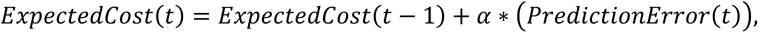

PE on each trial was computed as the difference between the expected cost (i.e., the likelihood of a task switch) and the experienced cost on trial, t (i.e., the actual task-switching likelihood of the effort level):

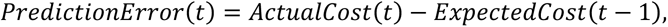

The rate at which the Expected cost is updated as a function of PE is controlled by a free parameter, *α*, which represents the learning rate of the participant. Given that our design did not include effort selections during the time of effort learning, we did not estimate learning rates based on a fit of effort selections. Previous simulation studies (Wilson & Niv, 2015) showed that model-based fMRI is insensitive to the change of learning rate parameters, as most experimental designs do not have the statistical power to detect differences in model parameters, specifically when the contrast-to-noise ratio and the number of trials is relatively low.

Consistent with this observation, we simulated learning rates between 0.2 and 0.7, in steps of 0.1 to test the effect of learning rates on the estimated BOLD responses of the SV Network ROI (Westbrook et al., 2019) for the expected cost regressor in Linear-Effort Level GLM model (see below for description of these models and ROIs). The results showed that there was no effect of learning rate parameters on the estimated BOLD response (Fig 2; *F*(1.01,18.24)=0.75, = .39, η_p_^2^ =.04). Consequently, we adopted a learning rate of 0.2 for the following analysis.

**Figure 2.**
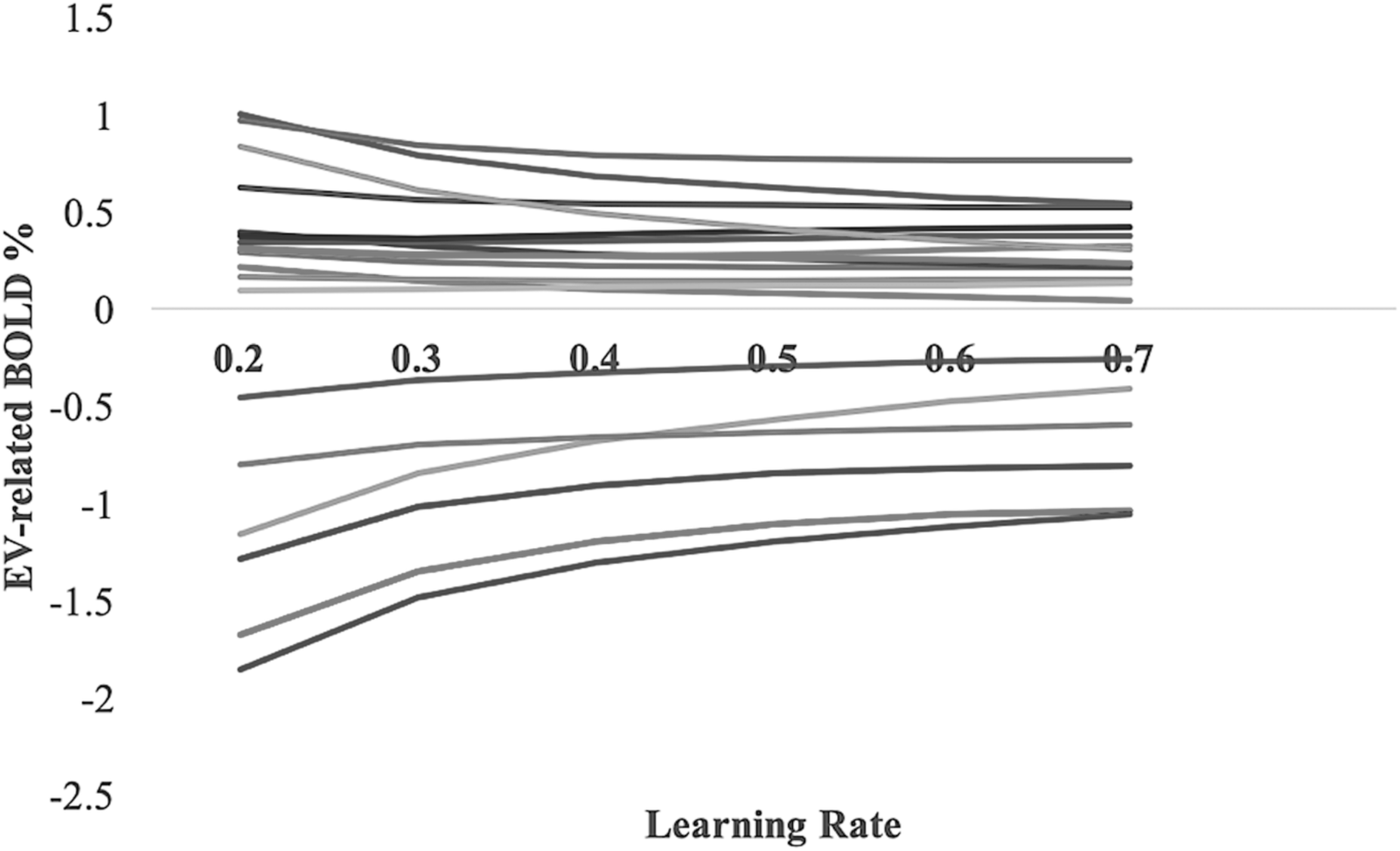
Regressor coefficients for expected cost regressor at the end of execution epoch estimated for SV Network ROI (Westbrook et al., 2019), as a function of learning rate. Each curve represents a single subject.

##### 2.3.1.2. Response Time Cost Models

The Response Time Cost Model learns the average time it takes to perform an effortful task based on context. We used two separate RT models that initialized the expected cost parameter in different ways.

First, we tested a model that initialized the expected cost at the average RT during the repeat trials of the first effort execution trial in the Learning Phase. Second, we tested a model that initialized expected cost as the first RT on the first trial of the first effort execution block in the Learning Phase. In other words, as we optimized effort execution epochs as blocks and not single-trial events, we calculated the average response time of each block as the ‘experienced cost’. Therefore, the ‘cost’ of each effort block (across 13 trials) is compared with the expected cost calculated across blocks of that effort level *so far*. Therefore, prediction error on the current trial was computed as the difference between the ‘experienced cost’ of the current effort block and the running average of all ‘costs’ across all execution blocks of that effort level.

Then, for both initializations, average RT across all trials of each effort execution block is calculated and the expected cost of each effort execution block is updated by a PE of that effort level with a learning rate of 0.2:

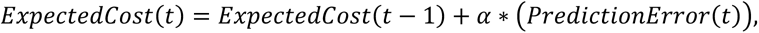

PE on each trial was computed as the difference between the expected cost (i.e., expected RT of that effort level so far) and the experienced cost on trial, t (i.e., the actual average response time in that effort run):

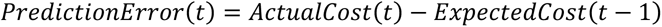

Next, in order to assess the influence of switch cost RTs in effort avoidance, we developed a separate model that assumes that the learned cost of effort is the experienced switch cost RTs in each effort block. This model is distinct from the RTcost model which assumes that the cost of cognitive effort is the average RT in each effort block, overall. Rather, the switch_cost_RT model only includes the difference between switch and repeat trials in a given effort block.

The initial expected cost (EV1) in the switch_cost_RT model was assumed to be the switch cost RT of the first effort block of the entire Learning Phase. In other words, we assumed that participants did not have an expected switch cost until they correctly switched for the first time in the Learning Phase. This initial switch cost was used as the initialization of EVs for all effort levels. Since effort blocks were pseudorandomized, all participants had the same effort level (level 3) as their first experienced effort block.

As Figure 4 shows, switch cost RTs linearly decrease with increasing effort level, while average RTs increase. The difference between the RTcost model and the switch_cost_RT model is that RTcost model assumes that effort is related to overall RT, and so the participant is considering the task as a whole, i.e., both switch and repeat trials, rather than only referencing the difficulty they experience on switch trials relative to repeat trials, which is the basis of the switch_cost_RT model. In other words, under an RTcost model, it doesn’t particularly matter for an effort cost whether an RT increase is due to a switch or some other factor. It is the average time taken to do trials of that task that matters, and it just so happens this is longer on average for tasks with lots of switching. As one key difference, on blocks with more task switching, repeat trials are known to take longer than the repeat trials of blocks with less switching (i.e., mixing costs). The RTCost model takes account of this mixing cost, as well, as a source of effort, whereas the switch_cost_model focuses only on the transition cost, the difference between switch and repeat.

**Figure 4.**
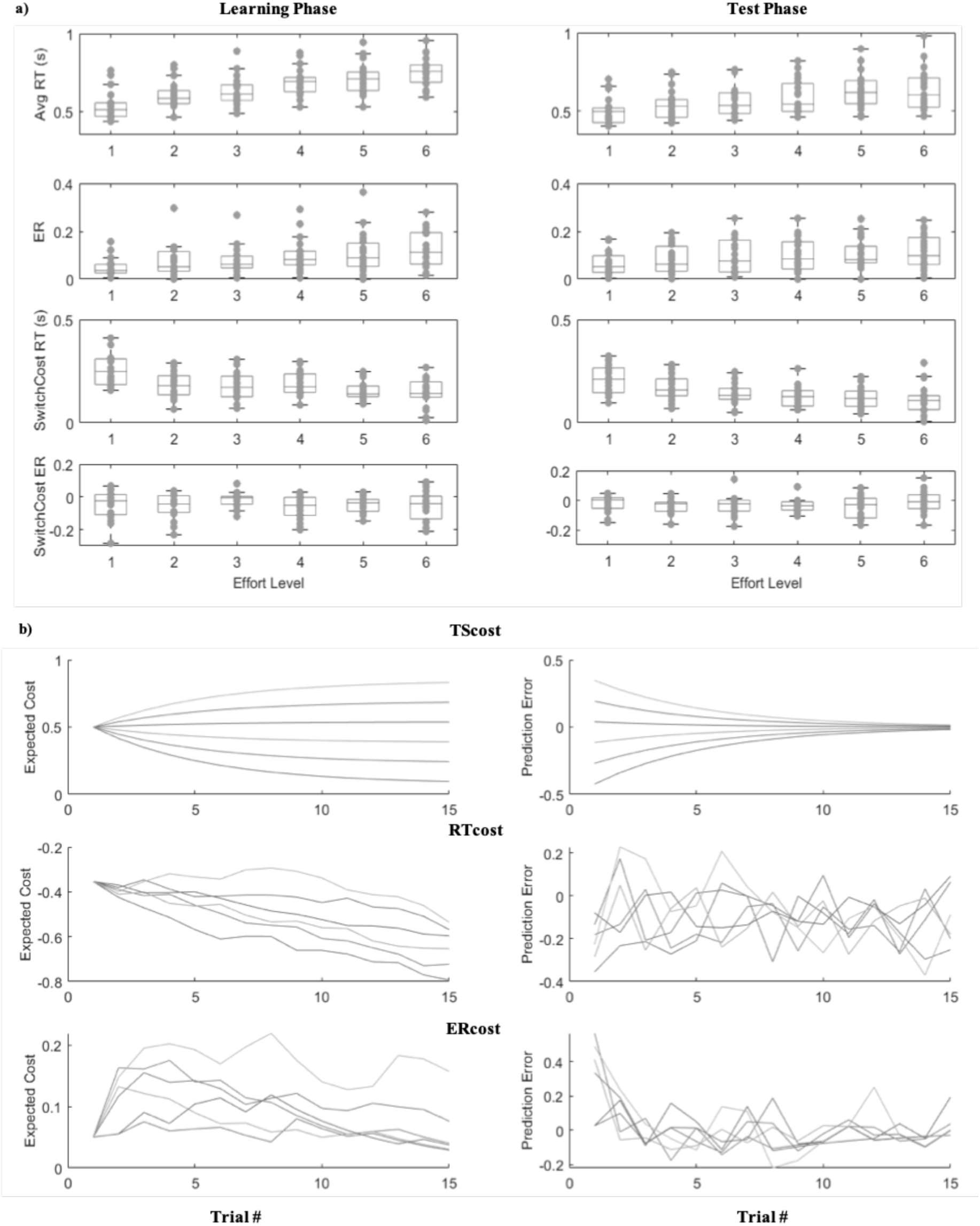
Task performance and model parameters. A) Average RT, ER, switch cost RT and switch cost ER for each effort level during the execution epochs of the Learning Phase and the Test Phase. Dots represent individual data points. B) Progression of expected Cost and Prediction Error model parameters across trials for TScost, RTcost and ERcost models. Each line represents a different effort level.

##### 2.3.1.3. Error Rate Cost Model

We additionally tested a separate model that assumed effort costs to be average error rates during effort execution during the Learning Phase. This model tests whether greater error-likelihood during the performance of difficult tasks is associated with an intrinsic cost with that task. Under this model, tasks with higher error-likelihood will be associated with less value during selection and will be avoided at higher rates during free choices of what task to do, all else being equal. In a trial-specific task-switching paradigm, we can use two measures of error likelihood: 1) average error likelihood of a task-switching block, 2) average switch cost error likelihood. The former measure involves the overall average error rate of all repeat, switch and initiation trials in the task block. In this model, expected cost was initialized at a 0.05 error rate and was updated based on the average error rate of a given effort run using the same Prediction Error algorithm explained above. The latter measure is the average error likelihood at switch trials minus the average error likelihood at repeat trials. In this model, we use the same initialization procedure used for switch_cost_RT model which assumes that the initial expected cost of all effort levels is the first experienced switch cost error rate at the first effort block. The latter measure has been shown to index the cognitive control cost associated with new task-set configurations (Rogers & Monsell, 1995). As such, in our analysis, we tested for the involvement of both measures.

### 2.4. MRI procedure

Whole-brain imaging was performed with a Siemens 3T Prisma MRI system using a 64-channel head coil. A high-resolution T1-weighted 3D multi-echo MPRAGE image was collected from each participant for anatomical visualization. Each of the two runs of the experimental task involved between 450 and 660 functional volumes depending on the participant’s response time, with a fat-saturated gradient-echo echo-planar sequence (TR = 2s, TE=28ms, flip angle = 90°, 38 interleaved axial slices, 192 mm FOV with voxel size of 3×3×3 mm). Head motion was restricted with padding, visual stimuli were rear projected and viewed with a mirror attached to the head coil.

#### 2.4.1. fMRI Analysis

Functional images were preprocessed in SPM12 (http://www.fil.ion.ucl.ac.uk/spm). Before preprocessing, data were inspected for artifacts and variance in global signal (tsdiffana, art_global, art_movie). Functional data were corrected for differences in slice acquisition timing by resampling slices to match the first slice. Next, functional data were realigned (corrected for motion) using 4^th^ degree B-spline interpolation and referenced to the mean functional image. Functional and structural images were normalized to Montreal Neurological Institute (MNI) stereotaxic space using affine transformation followed by a nonlinear warping based on a cosine basis set along with regularization, and then resampled into 2×2×2 mm voxels using 4^th^ degree B-spline. Lastly, images were spatially smoothed with an 8 mm full-width at half-maximum isotropic Gaussian kernel.

A temporal high-pass filter of 128 (.0078 Hz) was applied to our functional data in order to remove noise. Changes in MR signal were modeled under assumptions of the general linear model (GLM). Two GLMs were devised: a linear effort-level GLM and an independent effort-level GLM (described below). Both GLMs included nuisance regressors for the six motion parameters (x, y, z, pitch, roll, yaw) and four run regressors for the ‘Linear Effort-Level GLM’ and one run regressor for the ‘Independent Effort Level GLM’. The number of run regressors was different across GLMs because ‘Independent Effort GLM’ included the regressor for each effort level separately.

##### 2.4.1.1. Linear Effort-Level GLM

The linear effort-level GLM tested which voxels in the brain parametrically increased or decreased linearly with effort level during learning. Two event regressors were used. First, Execution events were modeled as an impulse function at the end of the presentation of the last trial stimulus of the sequence. Second, the Anticipation event regressor modeled each anticipation event with a fixed boxcar of 3 secs. We used orthogonalized parametric modulators on these event regressors to test the linear effect of effort level. The Execution event regressor was modulated in order by (a) a Prediction Error parametric regressor, and (b) an Effort Level parametric regressor corresponding to the effort level of that task sequence (1 through 6). As such, each Execution epoch was modeled as a block of all 13 trials that included both switch and repeat trials. PEs for each model were computed as the difference between the expected cost of the effort block minus the actual cost of the current effort block, as defined by each model (i.e., the running average ER, RT, average of switch cost RT or ER or the overall task-switching probability in that block). The Anticipation event regressor was modulated in order by (a) an expected cost regressor that scaled with the estimated expected cost of the upcoming effort execution (b) an Effort Level parametric regressor based on the presented effort level deck symbol (1 through 6). In order to test the effects of early vs. late learning on brain activity, the Execution and Anticipation event regressors, along with their parametric modulators, were modeled separately for each scanning run within the GLM. Two run regressors and a linear drift over the whole experiment were included as regressors of no interest.

##### 2.4.1.2. Independent Effort Level GLM

The independent effort level GLM sought to characterize the signal change related to each effort level independently of each other or of any particular function (e.g., linear). This GLM included twelve event regressors, one for each effort level (1 through 6) by epoch (Execution and Anticipation). Events in the Execution regressors were modeled as boxcars that onset with the presentation of the first trial stimulus of the sequence and ended with the participant’s response to the final item. Events in the Anticipation regressors were modeled with a 3 sec boxcar at the onset of deck symbol. Two run regressors and a linear drift over the whole experiment were included as regressors of no interest.

For both GLMs, SPM-generated regressors were created by convolving onset boxcars and parametric functions with the canonical hemodynamic response (HRF) function. To account for error due to differences in the HRF shape from the canonical, nuisance regressors were also created that convolved the onset functions with the temporal derivative of the HRF. Beta weights for each regressor were estimated in a first-level, subject-specific fixed-effects model. For group analysis, the subject-specific beta estimates were analyzed with subject treated as a random effect. At each voxel, a one-sample t-test against a contrast value of zero gave us our estimate of statistical reliability. For whole brain analysis, we corrected for multiple comparison using cluster correction, with a cluster forming threshold of *p* < .001 and an extent threshold, k, calculated with SPM to set a family-wise error cluster level corrected threshold of *p*<.05 for each contrast and group. Note that the higher cluster forming threshold helps avoid violation of parametric assumptions such that the rate of false positive is appropriate (Eklund et al., 2016; Flandin & Friston, 2016).

##### 2.4.1.3. ROI analysis

ROI definition is described below. For each ROI, a mean time course was extracted using the MarsBar toolbox (http://marsbar.sourceforge.net/). The GLM design was estimated against this mean time series, yielding parameter estimates (beta weights) for the entire ROI for each regressor in the design matrix.

We defined a fronto-parietal control (FPN) network ROI and Default Mode Network ROI derived from previously published cortical parcellations based on patterns of functional connectivity (Yeo et al., 2011). The FPN (Network 12 in Yeo et al. 2011) included bilateral prefrontal cortex, bilateral parietal cortex, and SMA. The DMN network (Network 16 from Yeo et al. 2011) included ventromedial prefrontal cortex, orbitofrontal cortex, posterior cingulate cortex and parts of precuneus.

As a complement to these network definitions, we also included definitions of FPN and DMN based on our prior work on effort avoidance. Our previous study (Sayali and Badre, 2019) parametrically manipulated implicitly learned effort cost/values in DST in fMRI. In that study, in order to have an unbiased analysis of the neural response function across effort levels, we conducted a Principal Component Analysis (PCA) of the whole brain to explore the shape of the neural functions of different brain areas that respond to effort execution during the Test Phase. Activity in these ROIs was associated in that study with effort avoidance decisions. To draw connection with that prior work, we also defined fronto-parietal control and DMN ROIs based on previous study’s Test Phase PCAs.

We additionally defined Subjective Value (SV) Network ROIs based on the regions cited by Westbrook et al. (2019) in order to test the involvement of those brain regions that have been shown to significantly track subjective value during effort discounting. Accordingly, we drew a 6 mm sphere around each region that significantly correlated with the subjective value of effort.

## 3. Results

### 3.1. Demand avoidance behavior

Overall, participants performed accurately during task execution across both phases (mean error: 9% in the Learning Phase, 9% in the Test Phase), and participants missed the deadline on few trials (2.6% of Learning phase trials, *SE*=0.01, 1.2% of Test phase trials, *SE*=0.03).

Task performance was impacted by the proportion of task switches across tasks. Across Learning and Test phases, there was a significant effect of effort level on both error rates and correct trial RTs (error rates: (*F*(5,85)=20.81, *p*< .001, η_p_^2^ =.56), RT: (*F*(2.03,36.62)=146.60, *p*< .001, η_p_^2^ =.90), and the effect of effort was linear for both (errors: *F*(1,17)=43.20, *p*< .001, η_p_^2^ =.72; RT: *F*(1,17)=214.49, *p*< .001, η_p_^2^ =.93). Additionally, correct trial RTs but not error rates reduced across Learning and Test phases (error rates: (*F*(1,17)=2.23, *p*> .05, η_p_^2^ =.01), RT: (*F*(1,17)=44.47, *p*< .001, η_p_^2^ =.72), indicating participants improved their speed but not accuracy across phases (Figure 4a).

We repeated the same analysis for switch cost RT and ER measures. Switch cost is a measure that takes the difference between switch trial RT/ERs and repeat trial RT/ERs. Across Learning and Test phases, there was a significant effect of effort level on switch cost error RTs but not switch cost ERs (switch cost RT: *F*(5,85)=18.72, *p*< .001, η_p_^2^ =.52; switch cost error rates: *F*(5,85)=1.37, *p*> .05, η_p_^2^ =.08), and the effect of effort was linear for switch cost RTs (*F*(1,17)=33.86, *p*< .001, η_p_^2^ =.67). Notably, this linear effect of switch cost across effort levels was negative, in that there was a reduced switch cost with more frequent switching. This is due to mixing costs which result in slower repeat trial RT in contexts with higher frequency task switching. Additionally, switch cost RTs but not switch cost error rates reduced across Learning and Test Phases (RT: *F*(1,17)=28.75, *p*< .001, η_p_^2^ =.63; error rates: *F*(1,17)=3.53, *p*> .05, η_p_^2^ =.17), indicating participants improved their speed in switching versus repeating trials but not accuracy across phases.

Effort level affected task selections, as expected. On average, participants avoided the harder task 67% (*SE*=0.02) of the time (Fig. 3a), which differed from chance (*t*(18)=6.87, *p*<0.001). The probability of avoiding the harder task did not change across Test blocks (*F*(3,54)=2.73, *p*=.53, η_p_^2^ =.13), indicating that participants’ effort selections were stable across time.

**Figure 3.**
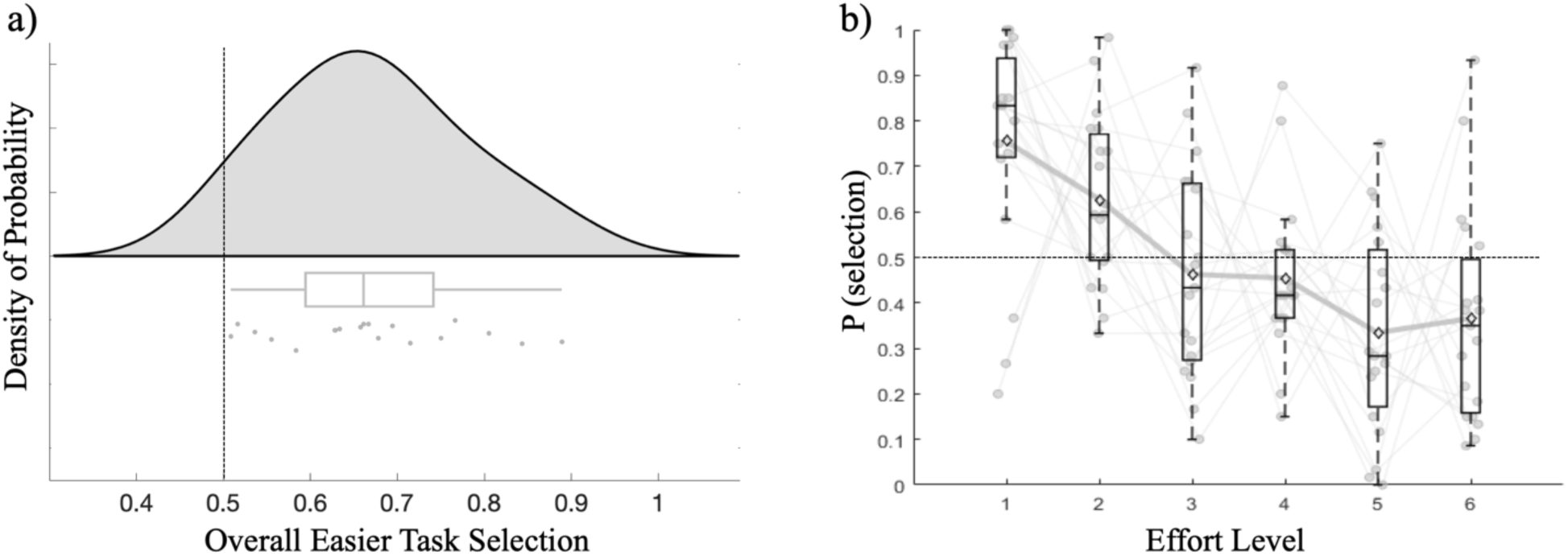
Demand avoidance across participants. A) Overall probability of selecting the easier task across all participants. The probability distribution of demand avoidance across subjects is plotted. Individual subjects are plotted as dots. All participants selected the easier task more than 50% of the time. B) Probability of selecting an effort level across participants. Dots represent individual data points. Open points represent the mean. The probability of selecting a task significantly decreased across all participants with increasing effort level. Light grey lines indicate chance (0.5) level.

Participants avoided higher effort levels at a greater rate than lower effort levels (*F*(5,90)=8.86, *p*<.001, η_p_^2^ =.33), and the rate of task avoidance was linear across effort levels (*F*(1,18)=47.88, *p*<.001, η_p_^2^ =.73; Fig. 3b), replicating Sayali and Badre (2019). The probability of selecting the easier task also significantly changed depending on the difference between effort levels for a given choice (*F*(2.30,41.35)=5.14, *p*=.001, η_p_^2^ =.22), such that larger differences increased the probability of choosing the easier task. This effect was also linear across effort levels (*F*(1,18)=13.48, *p*=.002, η_p_^2^ =.43). We observed a marginal increase in decision time across effort levels (*F*(5,85)=1.74, *p*=.06, η_p_^2^ =.09).

Consistent with the implicit nature of our effort learning paradigm, post-experimental debriefing inventory results showed that participants were mostly unaware of the difficulty manipulation across effort tasks. Across participants and decks, average ranking accuracy was 39%, or approximately 2 out of 6 rated correctly. Only 7 out of 19 participants reported that they noticed a difficulty difference between decks. Further, free text responses showed that even the “aware” participants were not completely sure about the exact difficulty ranking across decks. Across “aware” participants and decks, average rating accuracy was 57%, and only one “aware” participant (out of 19) was able to correctly rank all deck symbols.

### 3.2. Computational Modeling– Learning effects

We modeled effort learning using a reinforcement-learning model that acquires expectations about task performance (errors and RT) and task-switching probability for each task context (i.e., deck). In order to test our model estimates, we tested the effects of predicted response time, error rate and task-switching probability on subsequent effort selection behavior by conducting a mixed effects hierarchical regression analysis. Using a mixed-effect regression analysis, we tested which model explained selection rates better by testing the relationship between the finalized expected costs at each effort level during the Learning Phase and the selection rates in the Test Phase.

First, we tested the Error Rate Cost model that assumed effort costs to be the average error rates during effort execution in the Learning Phase. However, mixed-effects regression analysis showed that final expected error-rate costs did not explain selection rates during the subsequent Test phase, (t(114)=0.01, *p*=0.54). Next, we tested the Switch Cost Error Model that assumed effort costs to be the average switch cost error rate during effort execution in the Learning Phase. Mixed-effects regression analysis showed that final expected error-rate costs did not explain selection rates during the subsequent Test phase, (t(114)=-0.35, *p*=0.23) either. We repeated the same analysis by calculating expected values using a learning rate that increments with 0.1 steps between 0.2 and 0.7. None of the models revealed a significant relationship between expected error rate costs and selection rates, so we did not include the error rate cost model in further analysis.

Second, we tested the two separate RT models that differentially initialized the expected cost parameter. The first RT model initialized the expected cost as the average RT from the repeat trials of the first effort execution block. The second initialized expected cost as the first trial RT on the first effort execution block of the Learning Phase. The model that assumed initial expected cost to be the first RT significantly correlated with subsequent task selection rates during the Test phase (t(114)=-0.422, *p*< 0.001). This model performed almost the same as the model that assumed initial expected cost to be the average Repeat RT of the first effort run (t(114)=-0.418, *p*<0.001). Thus, we picked one of these models (first trial RT) for model-based fMRI analysis.

Third, in a hierarchical regression model, we observed that the final expected costs of the switch_cost_RT model did significantly explain effort avoidance rates (t(114)=0.48, *p*= 0.008). Though, here, greater switch cost RTs predicted a tendency to select a task rather than avoid it. This counterintuitive positive relationship between switch costs and task selections is likely due to increasing repeat trial RTs (i.e., mixing costs), which makes the transition switch cost smaller on blocks with more switching while overall RT goes up. Consistent with this, the effect of switch_cost_RT no longer explained effort avoidance rates in the presence of RTcost parameters (t(114)=0.32, *p*= 0.07), while RTcost still did (t(114)=-0.35, *p*= 0.003). These findings suggest that overall RT costs explain effort avoidance beyond the explanatory effect of only transition-based switch cost RTs. This is because RTcost model considers average correct RTs independent of trial type (e.g. switch or repeat), its cost parameter is influenced by any source of RT variance, such as post-slowing in RTs, switch costs, mixing costs and initiation costs. This makes RTcost model a comprehensive time-on-task model that accounts for both subjective control-related performance costs as well as opportunity cost of time. In other words, time-spent-on-task is a better predictor of effort avoidance than control related RTs.

Next, we tested whether task-switch probability explained selection rates. Note that the Task Switch Cost model updated expected costs by the difference between expected switch probabilities that are initiated at 0.5 and the actual switch probabilities of the effort task. Thus, given the trial numbers at learning, finalized expected costs always converged to the true switch probabilities of the effort tasks (see Table 1 for an example of average expected effort costs). Task-switch probability significantly predicted selection rates across participants, (t(114)=-6.83, *p*< 0.001).

**Table 1.** Finalized average expected costs values for RT cost models, Task Switch Cost model (TScostModel) and Error Rate cost (ERcostModel) models. RTcostModel1 refers to the model expected cost of the first trial is initialized at first trial of the first effort run of the Learning Phase. RTcostModel2 refers to the model expected of the first trial is initialized at the average repeat trial RT of the first effort run. Standard deviations (SD) are reported in parentheses. Note that for TScostModel, all finalized expected costs converged at the true task-switch probability of the effort block after 15 blocks, hence SD across participants was 0.

|  |  | Effort Level |  |  |  |  |  |
| --- | --- | --- | --- | --- | --- | --- | --- |
|  |  | 1 | 2 | 3 | 4 | 5 | 6 |
| Average<br>Finalized<br>Expected<br>Cost | RTcostModel1 | -0.70<br>(0.17) | -0.62<br>(0.15) | -0.56<br>(0.17) | -0.48<br>(0.14) | -0.44<br>(0.16) | -0.38<br>(0.14) |
|  | RTcostModel2 | -0.72<br>(0.17) | -0.64<br>(0.16) | -0.57<br>(0.18) | -0.49<br>(0.14) | -0.45<br>(0.16) | -0.39<br>(0.15) |
|  | ERcostModel | 0.05<br>(0.04) | 0.08<br>(0.07) | 0.11<br>(0.07) | 0.11<br>(0.08) | 0.12<br>(0.10) | 0.13<br>(0.09) |
|  | TScostModel | 0.08 | 0.23 | 0.38 | 0.54 | 0.69 | 0.85 |

Finally, we asked whether RT cost predicts selection rates over and above task-switch probability. In the presence of task-switching probability, neither RTcost model explained additional variance in selection rates (both *p*s>0.05). As the task-switch cost model alone explained 29% and RT cost models alone explained 10% of the variance in selection rates, model comparisons showed that including the RT cost expected costs in the same model did not significantly improve model fit (X2 (1, N = 2) = 0.4, *p* > .05). However, in order to understand the potential contribution of performance based effort learning and as RT cost does correlate with subsequent task selections, we included the RT cost model in our model-based fMRI analysis as well as the Task Switch Cost model.

Note that some participants’ effort selections deviated from the true ranked order in terms of probability of choosing an effort level across all other effort levels with which that effort task was paired. However, all participants almost always chose the easier effort task when paired with the harder effort task. For example, only 3 people out of 19 selected the harder task more than 50% of the time in a given effort pair. The performance model-based fMRI is agnostic to the true ranked order of the effort levels, as participants’ own performance on a current effort execution block determines the cost of that effort level. On the other hand, the task-switching cost model-based fMRI updates task value based on the objectively manipulated task-switch probability of the effort block. The latter suggests that for participants whose selections deviated from the true ranked effort order, expected cost, as defined by task-switch probability, might not fully account for their behavior. However, across all participants, task-switch probability explained 29% of the variability in selection behavior.

### 3.3. The functional form of univariate brain activity over effort levels

We next sought to replicate and extend our prior observations from the Test phase (Sayali and Badre, 2019) to the Learning phase in a new sample of participants. We investigated the pattern of fMRI activity across effort levels during the Learning Phase using ROIs defined from the Test phase of Sayali & Badre (2019, see Methods), as well as ROIs in the *a priori* SV network (Westbrook et al., 2019). Note that we also separately tested independent ROIs in FPN and DMN defined from a large scale study of resting state functional connectivity (Yeo et al., 2011). However, these ROIs showed similar trends to those reported below, so we do not detail them further.

As expected, activation in FPN showed a linear increase across effort levels during task execution (Fig. 5 left panel). There was a significant effect of effort on FPN activation (*F*(5,90)=11.12, *p*< .001, η_p_^2^ =.38) such that activity linearly increased with greater effort requirements of the task (*F*(1,18)=26.02, *p*< .001, η_p_^2^ =.59). Additionally, FPN activation decreased over time (in terms of experimental runs; *F*(1,18)=11.29, *p*= .003, η_p_^2^ =.39). There was no effect of anticipated effort on the FPN activation during the Anticipation epoch (Fig. 5 right panel: *F*(3.51,63.21)=1.22, *p*= .31, η_p_^2^ =.06).

**Figure 5.**
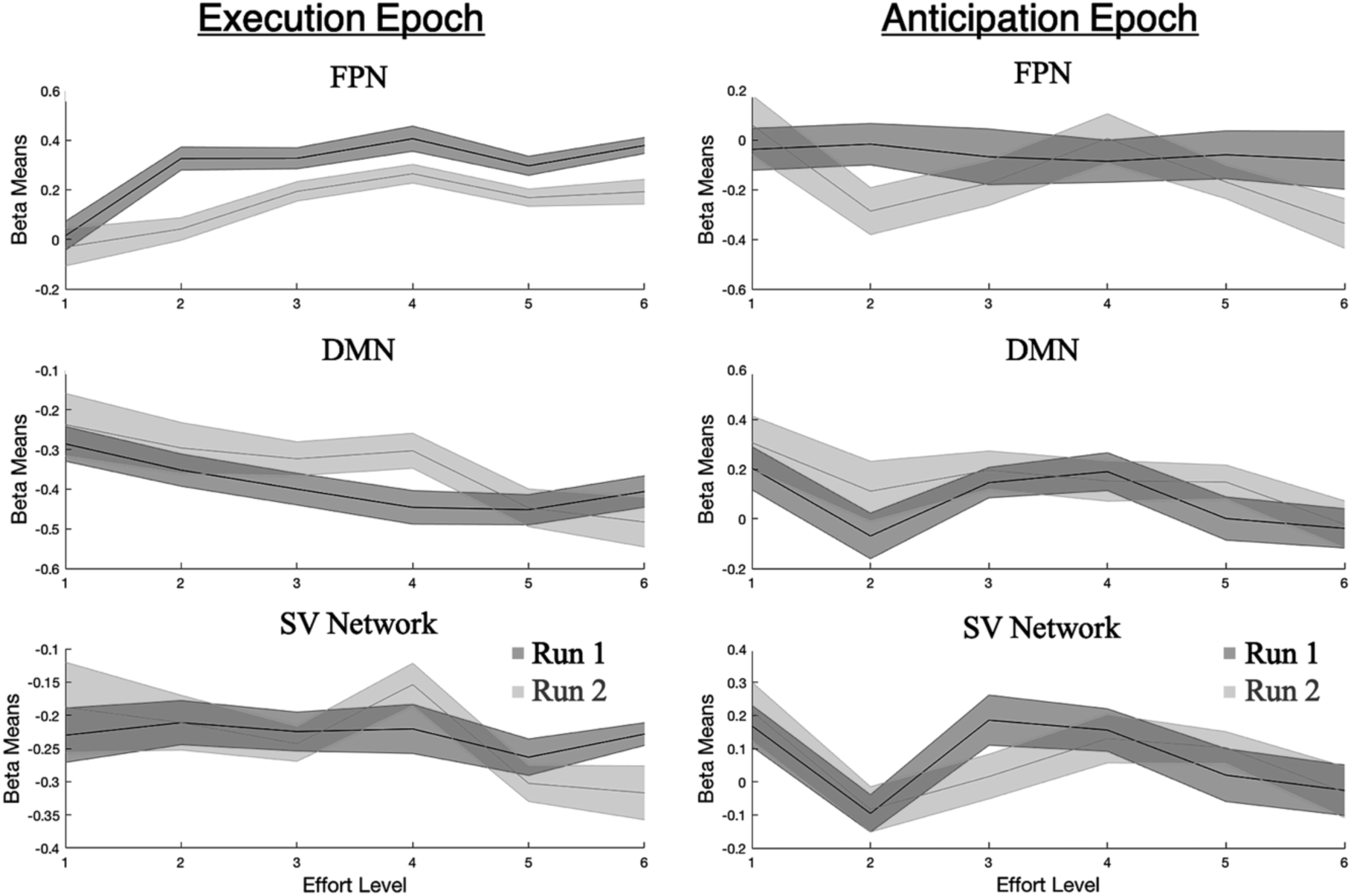
Regressor coefficients at the time of Execution and Anticipation Epochs for FPN component, DMN component and the Subjective Value Network. Different runs of the Learning Phase are plotted in different shades of grey in order to show the change of neural activity across time in the Learning Phase. Both FPN and DMN showed a linear trend across the execution of effort levels. FPN was the only ROI that changed its activity between runs during the Execution epoch. DMN showed a marginal reduction in activity across the anticipation of greater effort levels. All error bars stand for the standard error of the mean (SEM).

Consistent with our prior observation from the Test Phase (Sayali and Badre, 2019), during the Learning Phase, the DMN showed a negative linear trend in activation across effort levels (Fig. 5 left panel). There was a significant effect of effort on DMN activity (*F*(5,90)=3.31, *p*= .01, η_p_^2^ =.16) and a linear decrease in activity with increasing effort requirements of the task (*F*(1,18)=11.59, *p* = .003, η_p_^2^ =.39). There was no effect of time (in terms of runs) on DMN activity (*F*(1,18)=1.19, *p*= .29, η_p_^2^ =.06). There was a marginal effect of effort on the DMN during the Anticipation epoch (Fig. 5 right panel, *F*(5,90)=2.25, *p*= .06, η_p_^2^ =.11; Yeo et al definition of DMN: *F*(5,90)=2.34, *p*= .05, ηp2 =.12), suggesting that DMN marginally reduced activity in the anticipation of increasing effortful task exertion.

We did not find a significant positive or negative trend in SV network activity across levels of effort execution (*F*(5,90)=1.23, *p*= .30, η_p_^2^ =.06) or across blocks during task execution (*F*(1,18)=0.03, *p*= .26, η_p_^2^ =.002). During the anticipation interval prior to each imposed task, SV network activity was positively related to effort level (*F*(5,90)=2.44, *p*= .04, η_p_^2^ =.12, Fig. 5 right panel). Within-subject contrast results showed that the SV network showed a cubic trend across the anticipation of effort levels (*F*(1,18)=5.70, *p* = .028, η_p_^2^ =.24). There was no effect of run during the anticipation epoch on activity in SV Network (*F*(1,18)=0.02, *p* = .88, η_p_^2^ =.001).

In sum, the FPN and DMN exhibited similar modulation of brain activity during the Learning of effort tasks as previously observed during the Test phase. Further, FPN recruitment decreases with increased experience in the Learning Phase, which is expected based on the prior literature on task learning and experience (Ruge and Wolfensteller, 2010; Bhandari and Badre, 2020). The *a priori* SV Network showed a reliable omnibus effect of increase activity with greater anticipated effort prior to task performance.

### 3.4. Model-based fMRI

#### 3.4.1. Task Switch Cost model

Our modeling analysis of learning behavior indicated that the expected likelihood of task switching acquired during the Learning phase was the best predictor of subsequent task selections during the Test phase. In the model, this expected likelihood is formed by incrementally adjusting an expected task switch probability based on task switch prediction errors. We next sought to test how the *a priori* hypothesized networks, FPN, DMN, and SV network, track prediction errors related to task switch probability. Thus, we tested the contribution to activity in these brain regions from the parametric modulation of Prediction Error and expected cost parametric regressors in these ROIs, in addition to the Execution Level and the onset of Execution and Anticipation events. We estimated brain activity for each experimental run of the Learning Phase separately in order to test the effects of early vs. late effort learning on brain activity. These effects were modeled within the Linear Effort Level GLM, described in the Methods.

We focus here on the key model-based results related to prediction error and expected cost (see Fig. 6). The FPN correlated positively with prediction error (*p* < 0.001), such that FPN activation was greater when tasks switched more frequently during a block than expected based on prior experience (Fig. 6). This effect was strongest during the second relative to the first run of the experiment (t(18)=-4.56, *p*<0.001). Neither the DMN nor the SV network correlated with prediction error (both *p*s > 0.05), despite strong trends, particularly for the former. Neither FPN, DMN or SV networks correlated with expected costs.

**Figure 6.**
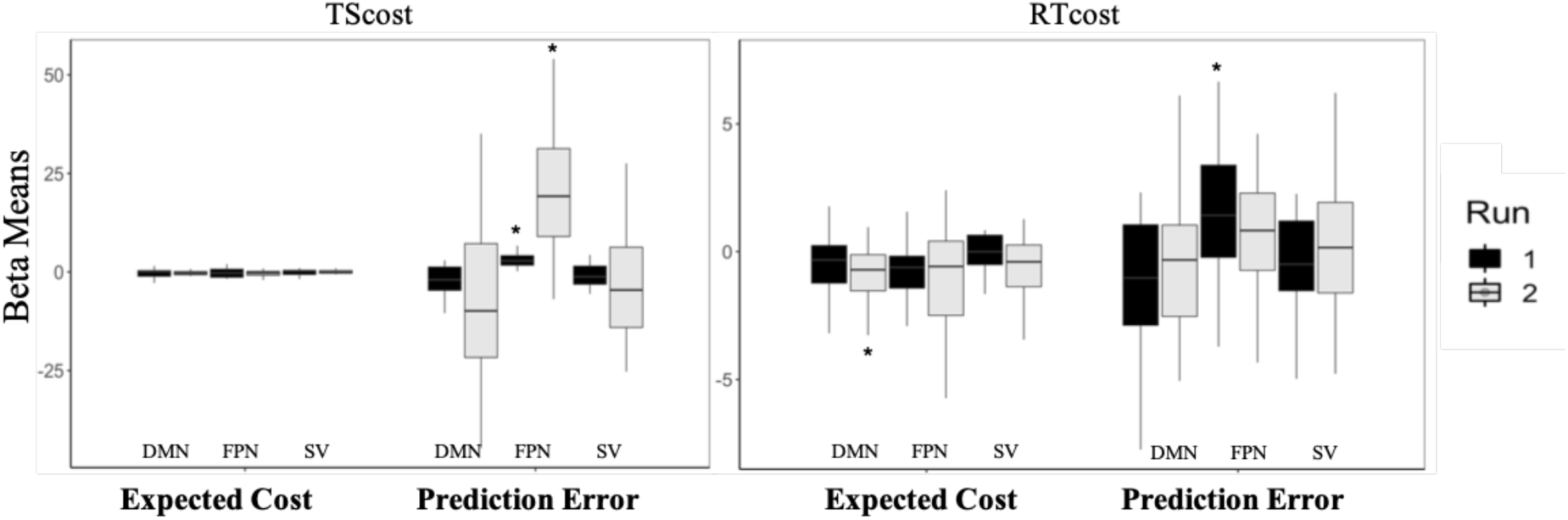
Regressor coefficients for Linear Effort Level GLM for DMN, FPN and Value Network ROIs. FPN positively scaled with PE during both experimental blocks for both Task Switch Cost Model (TS_Cost) and during the first block for Response Time Cost Model (RT_Cost). PE and Expected Cost parameters are parametric regressors associated with the Execution and Anticipation epoch onset regressors respectively.

In order to test specific nodes within the hypothesized SV network, we analyzed additional ROIs in left, right VS, two rVMPFC ROIs, lVMPFC and dACC (Supplementary fig. 2) that were included in Westbrook et al., (2019). While both left and right vMPFC showed significant deactivations during effort execution and anticipation, only dACC (Figure 7) significantly tracked PE regressor during the first run (β=3.50, p< 0.001) and parametrically increased with effort anticipation in the second run (β=1.57, p< 0.001).

**Figure 7.**
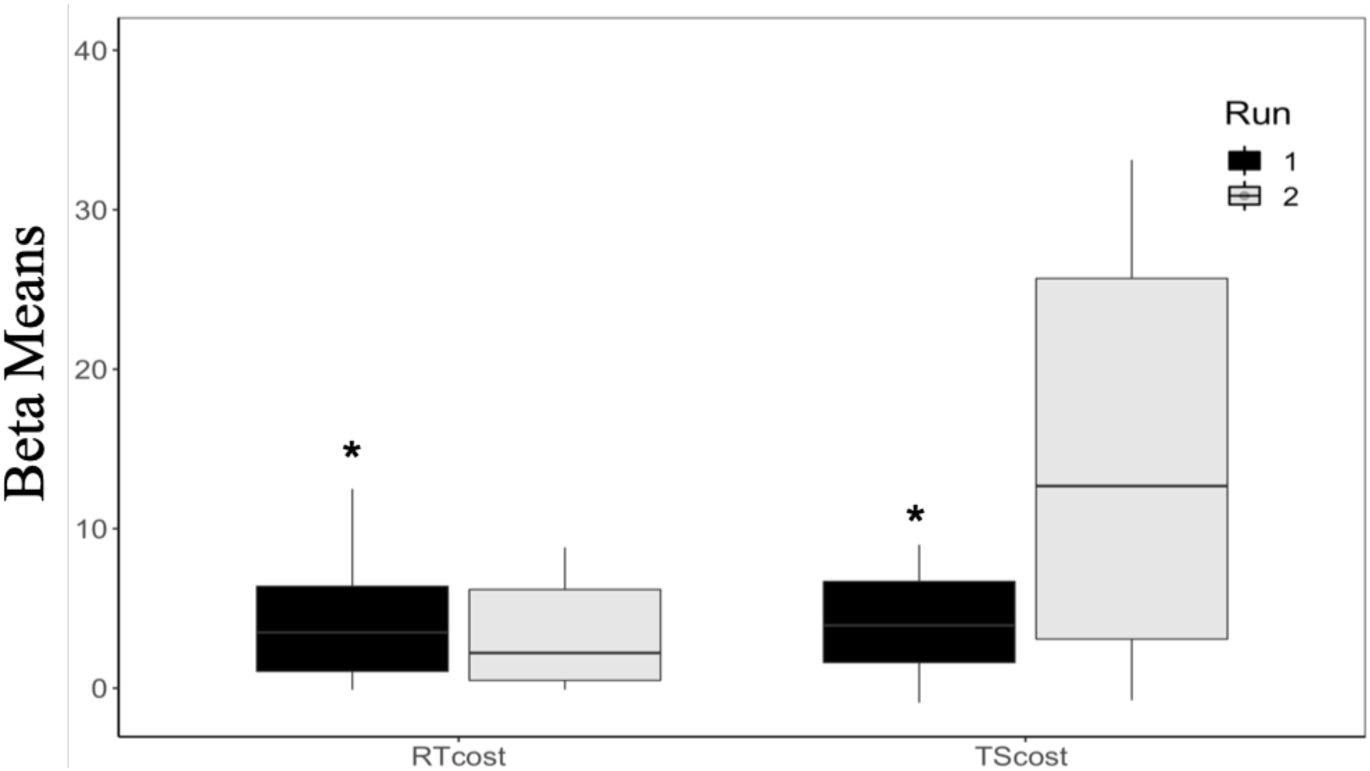
Average dACC activity for RTcost and TScost model prediction error parameters. For both models, prediction error significantly correlated with dACC activity.

Finally, to complement the ROI analyses, we performed a whole-brain voxel-wise analysis (Fig. 8) of the key model-based regressors. As shown in Figure 8, the expected cost regressor negatively correlated with one cluster at left middle frontal gyrus (x=-26, y=4, z=46). Two clusters positively correlated with prediction error (Figure 8). One cluster was centered at superior parietal lobule (x=-12, y=-68, z=56) and the other was centered at left middle frontal gyrus (x=-40, y=24, z=24).

**Figure 8.**
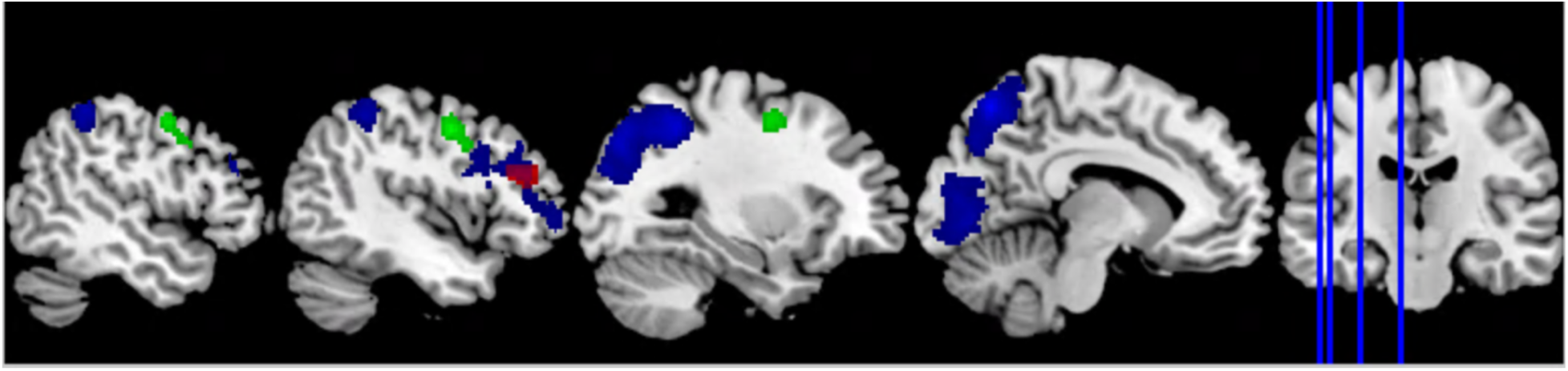
Whole Brain Results for Task Switch Probability Cost and Response Time Cost Models. All results thresholded at FWE p < .05. **Figure legend**: TScost model PE+ printed in blue; RTcost model PE+ printed in red; TScost model Expected Cost-printed in green.

#### 3.4.2. RT cost model

As the RT cost model also predicted selection rates, we repeated our fMRI analysis using expected values and prediction errors derived from this model (Fig. 6).

The whole brain analysis based on RTcost PE yielded similar results with that of Task Switch Cost PE. One cluster survived the family-wise error-correction (Figure 8). Peaks positively correlated with PE were identified in left middle frontal gyrus (x=-42, y=32, z=24).

ROI results showed that the FPN correlated with prediction error in the 1^st^ run but not the 2^nd^ run, though these correlations were not significantly different. Given that TScost model increased its correlation by the second run whereas the RTCost model did not, there was a reliable model*run interaction (*F*(1,57)=21.91, *p*<0.001). Neither the DMN or the SV network correlated with prediction errors (both *p*s > 0.05), and neither FPN or SV networks correlated with expected costs. Importantly, Expected Cost regressor negatively correlated with DMN during the 1^st^ run of the Learning Phase (p<0.01; Fig. 6). This relationship persisted even after SV regions that overlap with DMN (such as in VMPFC and PCC) has been removed, indicating that DMN but not SV predicted expected effort costs as tracked by time-on-task.

## 4. Discussion

A central problem in cognitive neuroscience has concerned why people choose to perform certain tasks over others when prospective rewards are otherwise comparable. The widespread view is that cognitive tasks are subjectively costly and discount outcomes accordingly. Thus, an open question has been what aspects of a task make it costly and how does the brain track these demands. The present study adds a new perspective on this question by considering which demands are tracked during experience with a task, in the absence of any decision, but that later correlate with avoidance.

We find evidence that at least two factors during effort learning explain later effort avoidance rates: the frequency of task switching and the expected time-on-task. These factors are correlated, and so it was hard to distinguish them behaviorally. Indeed, task switching frequency held a stronger relationship to choice and was able to fully explain the behavioral variance that was accounted for by expected time-on-task. However, this observation does not conclusively rule out other contributors. As such, we tested the effect of RT-related costs on effort learning in the brain using model-based fMRI.

Taking this approach, we found that trial-to-trial PE from each model held stronger relationships with the FPN at different stages of learning. Specifically, when RT was greater than expected, PE-related brain activity in FPN control networks was strong both early and late in learning. In contrast, the correlation of FPN brain activity with PE derived from the expected frequency of task switching was primarily in the second run of learning.

This observation is intriguing, but we note that we did not hypothesize these differences a priori. Instead, we had predicted that the SV network would correlate with one or both of these measures of task demand; a prediction for which we found limited evidence and that we discuss below. Nonetheless, if verified in subsequent work, these observations offer insights into the nature of task demand learning. At the surface, they suggest that multiple factors likely contribute separately to our assessment of task demands. Thus, rather than a single demand that drives all evaluations of cognitive effort, multiple factors are weighed, likely dynamically, depending on the situation. For example, there may be contexts where time-on-task is more or less salient than the frequency of task switching. And further, there may be more than one brain system that contributes to these decisions.

The early association of control network activity with PE derived from RT rather than task switch frequency is particularly intriguing in light of the broader literature on task learning and automaticity. It is well established that when new tasks are performed, the first trial is associated with a higher RT and error rate that declines in log form toward an asymptote (Bhandari and Badre, 2018). Studies testing these early trial effects have repeatedly associated this period of rapid task adjustment with activation in control networks (Cole et al., 2010; Ruge and Wolfensteller, 2010; Mohr et al., 2016; Hampshire et al., 2019). A recent study that held the stimulus-response rules constant, but required adjustments only in control settings – analogous to the task switching demands that distinguish context in the present study – specifically found these rapid changes to be tied to adjustments in control settings (Bhandari and Badre, 2018). Following from this discussion, one hypothesis is that early learning is characterized by adjustments in control settings that adapt to the specific task context. If optimization at this stage is closely tied to changes in RT, then PEs related to RT are more informative regarding task demands.

The later sensitivity of the FPN to task switching frequency might similarly be related to the nature of task learning. The association of FPN with task switching has been widely observed (Kim et al., 2012). We also previously observed increasing activation in FPN with task switching frequency (Sayali and Badre, 2019), an effect we replicate here. However, the present results further show that the FPN is also more active when task switching is greater than expected. The later emergence of this association is notable given the overall activation decrease in the FPN with greater task experience.

When people learn control settings, in addition to the fast adjustments in RT noted above, there is also a slower process of learning that favors stable performance over variance in task demands (Bhandari and Badre, 2020). It is possible that the engagement of control, and the FPN, in this stable phase is attuned to unexpected and irregular events, particularly when they require control. Thus, deviations in the use of control itself, such as caused by task switching, can drive learning about task demands in this later phase.

The dACC may also be important in tracking these errors of prediction, particularly early in the task. We observed that dACC correlated with PE derived from response time and expected task switching frequency early in learning. This interpretation is broadly consistent with current views that ascribe a monitoring role to the dACC (Shenhav et al., 2013; Botvinick, 2007). As one example, the Predicted Response-Outcome Model (PRO; Alexander and Brown, 2011) argues that the medial portion of PFC, including the dACC, implements two functions: 1) learning to predict response outcome requirements, 2) signaling prediction errors. Accordingly, dACC activity reflects changes in the predicted value of the effortful task.

An important and presently open question concerns why our system focuses on learning about factors like time on task or task switching frequency as the basis for cognitive effort costs in later decision making. Though the present study does not directly address this question, our results are in line with at least two previously proposed accounts: 1) opportunity cost and 2) learning progress motivation.

The opportunity cost account proposes that we penalize certain tasks with effort costs because performing these tasks prevents us from gaining value from other tasks we could be performing instead or in addition (Kurzban, 2013; Musslick et al., 2018). For example, Kurzban (2013) argued that since we have limited ability to multitask, we penalize tasks that do not permit simultaneity and so the ability to acquire value from other tasks. As these alternative tasks and their subjective value might be impossible to determine, an extension of this account suggests that average reward rate of the environment as a function of time-spent-on-task is what updates task value (Niv et al., 2007). A similar idea might relate to the average stability or switch rate in the environment. Thus, the longer we take on a task or the more we use general purpose control machinery for task switching the more a task is penalized because these accordingly prevent us performing other tasks and result in an opportunity cost.

A mutually compatible account focuses on learning progress as the source of motivation, including effort costs. This account suggests that learnability of a task contributes to its long-run subjective cost (Gottlieb & Oudeyer, 2018). Accordingly, one might prefer those tasks on which one most improves task performance (e.g. reduce prediction errors) to those that improve slowly. This might be related to the emphasis the opportunity cost model places on simultaneity, in that automatic tasks can often be shared more easily with other tasks. So, in a sense, if a task appears more learnable, then regardless of its difficulty now, its learnability might allow one to discount future opportunity costs by practicing it. From the perspective of this account, our results might suggest that early impressions of learning, that are characterized by rapid changes in RT, may be crucial for assigning effort costs.

An important, distinguishing point to emphasize regarding the present task is that the participants were not explicitly aware of the difficulty difference between decks. We view learning in this task, then, as analogous to non-declarative learning wherein people acquire knowledge about regularities in their world that they can exhibit in their behavior, in the absence of conscious awareness (e.g., Gluck et al., 1988; Knowlton et al., 1994). These effects of learning without awareness have been observed in a wide range of contexts, including perceptual learning, statistical learning, category learning, rule learning, priming, and so forth. And indeed, many of these studies, focus on the reinforcement learning and value-based system that we test here as the basis for this kind of learning in the brain (e.g., Knowlton et al. 1994). Akin to these studies, in our paradigm, participants received no explicit instructions about what different decks of cards represented or on what basis they should make their selections. And, like many non-declarative learning studies (Gluck, Shohamy & Meyers, 2002; Knowlton et al., 1996; Squire, 1994), our participants reported no explicit knowledge of what they had learned during the debriefing phase. Given the probabilistic nature of the task-switching blocks, we assume that participants implicitly learned the statistical task structure, which then informed their decision making. Though there was no explicit feedback, participants did experience effort, which our model assumes they interpret as a cost for reinforcement learning.

We note that the implicit nature of the learning in our study, though analogous to other kinds of non-declarative learning, is different from other procedures for testing effort in the literature. Previous studies have shown that people avoid cognitive effort when they are explicitly instructed about differences in the difficulty of various options. Further, in some cases, people’s ability to explicitly detect effort difference between options was associated with greater effort avoidance (Naccache et al., 2005; Gold et al., 2015). For example, schizophrenia patients, compared to controls, showed difficulty detecting effort differences between options despite experiencing greater switch costs in a DST paradigm, suggesting that successful monitoring of effort costs rather than their implicit experience might underlie effort avoidance. Thus, future studies should distinguish what role explicit awareness plays in the learning of effort costs.

Based on previous studies of effort-based decisions and learning (Westbrook et al., 2019; Nagase et al., 2018), we hypothesized that the SV network would also track PE during the implicit learning phase. Surprisingly, we did not locate evidence of an association of either PE or anticipated cost with activation in the SV network. This seems at odds with prior observations, though at least one prior study also failed to locate an association of the SV network with the degree of physical effort required by a task (Prévost et al., 2010). However, given some effects in specific regions, such as those in dACC, it is possible that our study simply lacked sensitivity in this network to locate an effect over noise.

Another possibility is that the SV network is affected by the requirement to make a decision. A clue in this regard comes from the anticipation phase, when we would predict less activation in SV prior to a difficult task. Instead, we observed greater activation in the SV network during anticipation of a more effortful task. This reversal might reflect that participants are anticipating an imposed task rather than evaluating which task to perform. Indeed, Schouppe et al. (2014) reported that the direction of activation in VS during the anticipation of effortful tasks changed as a function of whether the task was imposed or voluntarily selected.

Instead of the SV network, we did find that that anticipated costs, based on time on task, tracked by the activity of the DMN during the cue epoch of the upcoming imposed effort task. Importantly, our neural definition of DMN comes from our previous effort avoidance study (Sayali & Badre, 2019), in which we found that an activity change in this ROI across effort levels predicted effort avoidance during decision-making across subjects. Thus, the present results expand the observation of cost-related activity in this network to the anticipation of costs for an upcoming task during learning. Given this observation across two studies, future work should more directly investigate the role of the DMN in effort cost learning and decision making.

Finally, we highlight three limitations of the study. First, as participants did not make task selections during learning, it was not possible to estimate individual learning rates through model fitting. This shortcoming indicates that current study cannot estimate the degree to which task switch cost or RT cost during effort learning is directly tied to effort selections. This will be an important gap to fill in future work.

Second, the present study prospectively sought to enroll and test demand avoiding participants. The rationale for this choice was that our previous study found significant individual differences in demand avoidance behavior on the basis of those who are demand seeking versus demand avoiding (Sayali and Badre, 2019), with demand seeking participants showing diminished or even reversed task choice behavior. We further found that participants in fMRI experiments tend to be disproportionately demand seeking. Thus, as the present study, was focused on understanding the neural basis of how effort costs are learned such that they lead to later avoidance, we used the NfC inventory to pre-screen our participants in order to enroll participants why were effort avoiding. Therefore, we used an independent measure to preemptively select our fMRI volunteers to test only those individuals for whom effort might register as a cost and not as a reward. Though this approach successfully decreased the individual variance in our study, this approach necessarily affects the generalizability of these results. Thus, our conclusions are most closely generalizable to demand avoiding individuals. Nonetheless, we argue that these effects are likely also applicable to less avoidant individuals when they choose to avoid effort. Indeed, in our prior work, we found that brain systems highlighted here, showed similar activity patterns across both demand avoiders and demand seekers as a function of task demands. However, it was the interpretation of these signals, in terms of their relationship to choice, that distinguished the groups.

Third, the current study tested a range of different hypothesized cost signals in separate models in order to distinguish which best explained behavioral and brain responses. As such, across models, we assumed that participants learned the cost associated with response times, accuracy performance, or objective task switching probability separately. However, we acknowledge that there could be interactions between these different costs that are not considered here. One example, would be to integrate speed and accuracy, such as implementing a drift diffusion approach or an Δ_accuracy_ variable that updates the current trial’s accuracy, in relation to the participant’s overall response time. However, given the number of trials each effort block yielded at each accuracy level, it was not possible to take such an approach in the present study. Future studies should explore whether speed and accuracy together affect effort costs or whether other combinations of task-related factors have a non-linear impact on learned effort costs.

What does experience with a task teach us about its demands? Our results indicate that at least two factors are critical. First, response time is tracked as an index of performance, particularly during our first experiences with a task, when we are rapidly adjusting our control settings and automatizing performance. We form expectations about those control settings over time, like task switching frequency, such that later deviations from these expectations become a prime source of learning about task demands. These expectations, the way they are modeled in the current study, may be generalized into real-life settings.

The results of the task-switch probability model can be generalized to any task that requires multi-tasking in a serial manner. For example, switching between checking your email and writing a manuscript or switching among apps on your smart phone will incur a switch cost in both time and quality of work. Further, in contexts with greater frequency of switching, the overall performance costs will be greater. Thus, over time, learning the switch frequencies associated with a particular context might shape future selection behavior. (As one personal example, one of us has come to dread a particular day of the week because it holds many meetings and other tasks both professional and personal, and so it is associated with lots of task switching throughout the day). As such, one might learn to avoid planning days with lots of switching to reduce these costs. Similarly, the results of the response time cost model might hold implications for any cognitive task that benefits from practice, as this performance cost is not specific to a task-switching paradigm. Our results suggest that first impressions of time-on-task may be more critical than later time costs in our effort estimations.

Importantly, however, our results indicate that no single factor likely drives effort costs. Future experimental work directed at effort learning will be important to determine what factors drive effort costs and in what contexts we learn about these costs for different kinds of tasks.

None of the experiments was preregistered. Data sets are available upon request to the corresponding author.

## Acknowledgements

This work was supported by a MURI from the Office of Naval Research (N00014-16-1-2832) and a grant from James S. McDonnell Foundation. We would like to thank Louise Stolz, Jordan Rubin-McGregor and Adriane Spiro for assisting in data collection, and Eliana Vassena for her helpful comments.

## Supplementary Material

**Supplemental Figure 1.**
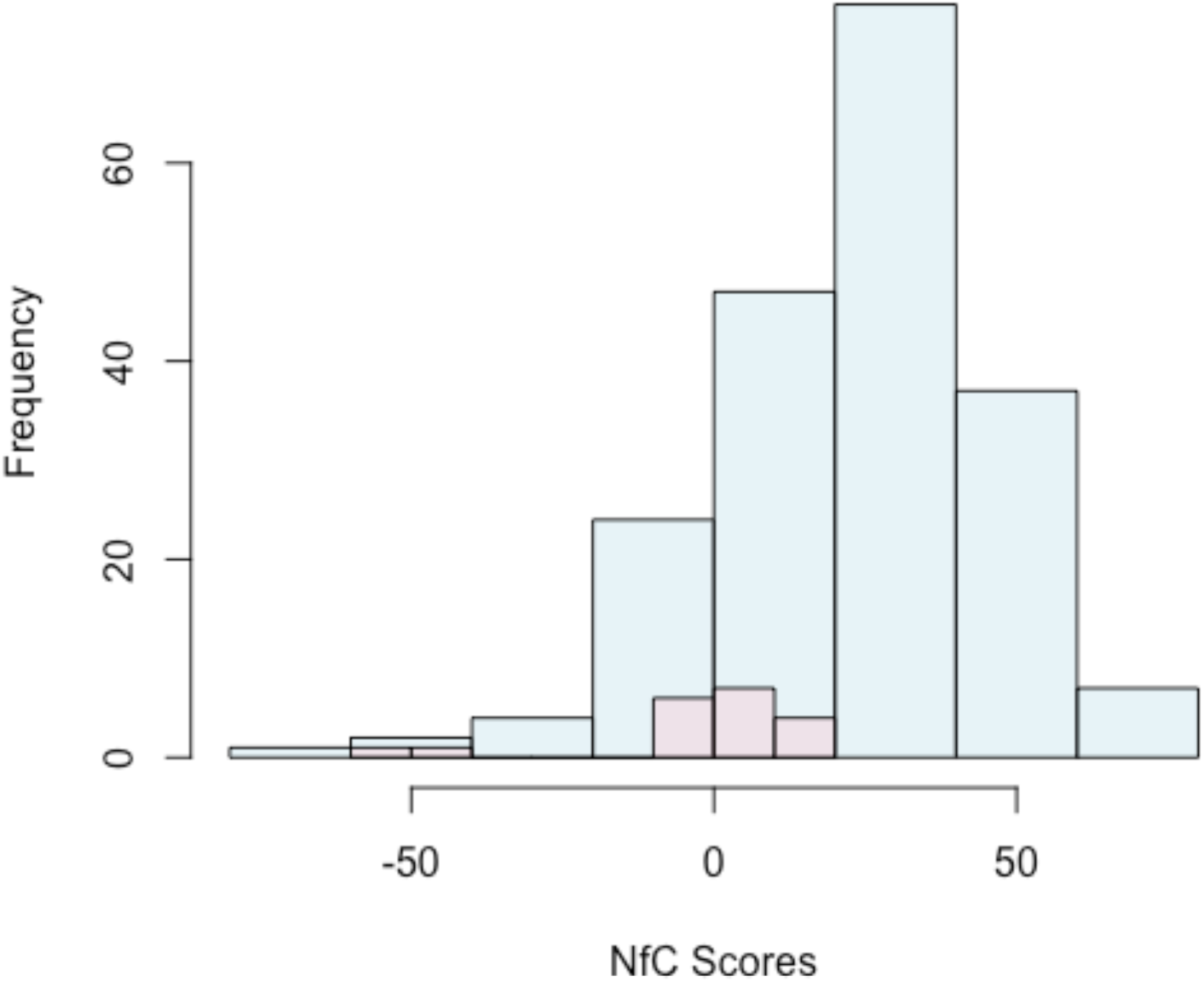
Histogram of Need for Cognition (NFC) pre-screened participants and fMRI volunteers. fMRI eligible participants were pre-screened with NfC inventory (bars in blue). 19 volunteers who scored lower than 16 (bars in red) on NfC were included in the study.

**Supplementary Figure 2.**
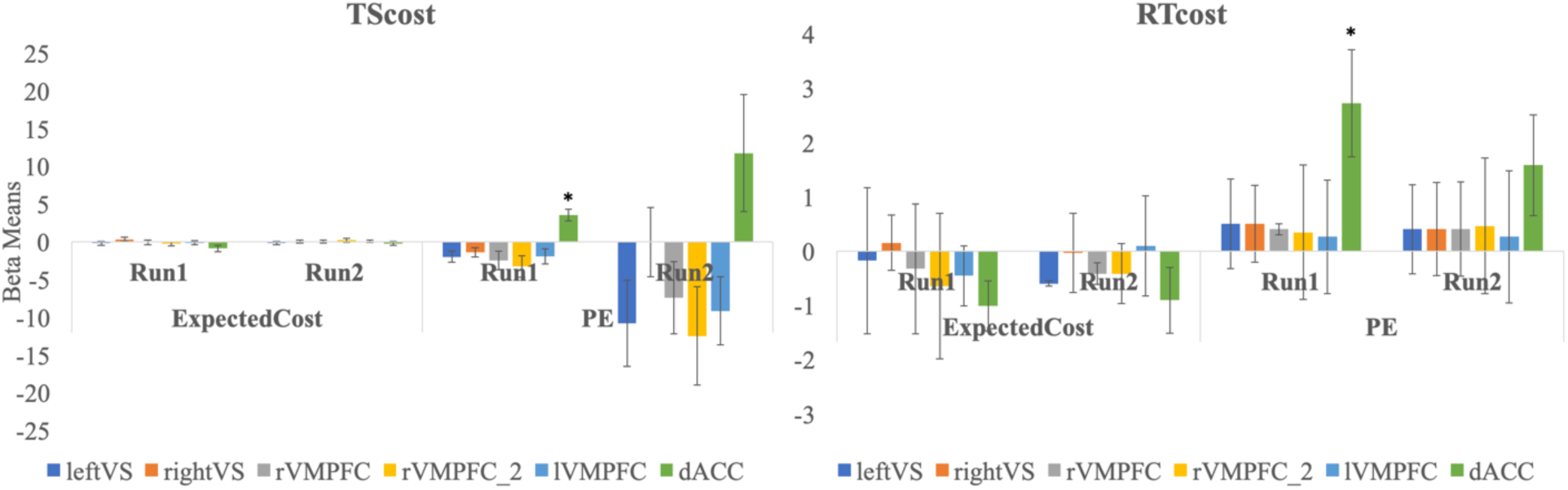
Beta weights for TScost and RTcost model parameters for Subjective Value Network ROI nodes. dACC positively correlated with prediction errors of both models in the first run of the Learning Phase.

